# Effects of tDCS Dose and Electrode Montage on regional cerebral blood flow and motor behavior

**DOI:** 10.1101/2021.02.02.429369

**Authors:** Anant Shinde, Karl Lerud, Fanny Munsch, David C Alsop, Gottfried Schlaug

## Abstract

We used three dose levels (Sham, 2mA, and 4mA) and two different electrode montages (unihemispheric or bihemispheric) to examine DOSE and MONTAGE effects on regional cerebral blood flow (rCBF) as a surrogate marker of neural activity, and on a finger sequence task, as a surrogate behavioral measure drawing on brain regions targeted by transcranial direct current stimulation (tDCS). We placed the anodal electrode over the right motor region (C4) while the cathodal or return electrode was placed either over a left supraorbital region (unihemispheric montage) or over the left motor region (C3 in the bihemispheric montage). Performance changes in the finger sequence task for both hands (left hand: p = 0.0026, and right hand: p = 0.0002) showed a linear tDCS dose response but no montage effect. rCBF in the right hemispheric perirolandic area increased with dose under the anodal electrode (p = 0.027). In contrast, in the perirolandic ROI in the left hemisphere, rCBF showed a trend to increase with dose (p = 0.053) and a significant effect of montage (p = 0.00004). The bihemispheric montage showed additional rCBF increases in frontomesial regions in the 4mA condition but not in the 2mA condition. Furthermore, we found correlations between rCBF changes in the right perirolandic region and improvements in the finger sequence task performance (FSP) for the left and right hand. Our data support not only a strong direct tDCS dose effect for rCBF and FSP as surrogate measures of targeted brain regions but also indirect effects on rCBF in functionally connected regions (e.g., frontomesial regions), particularly in the higher dose condition and on FSP of the ipsilateral hand (to the anodal electrode). At a higher dose and irrespective of polarity, a wider network of sensorimotor regions is positively affected by tDCS.

**Highlights:**

1. tDCS-DOSE had a linear effect on finger sequence performance for both hands
2. rCBF changes in both perirolandic ROIs demonstrated tDCS-DOSE effects, and left perirolandic ROI demonstrated tDCS-MONTAGE effects.
3. Simulated current intensity in the left and right perirolandic ROI strongly correlated with the contralateral hand’s finger sequence performance.
4. tDCS-Tolerability scores did not correlate with change in rCBF or finger sequence performance of the left hand.

**Graphical Abstract:** 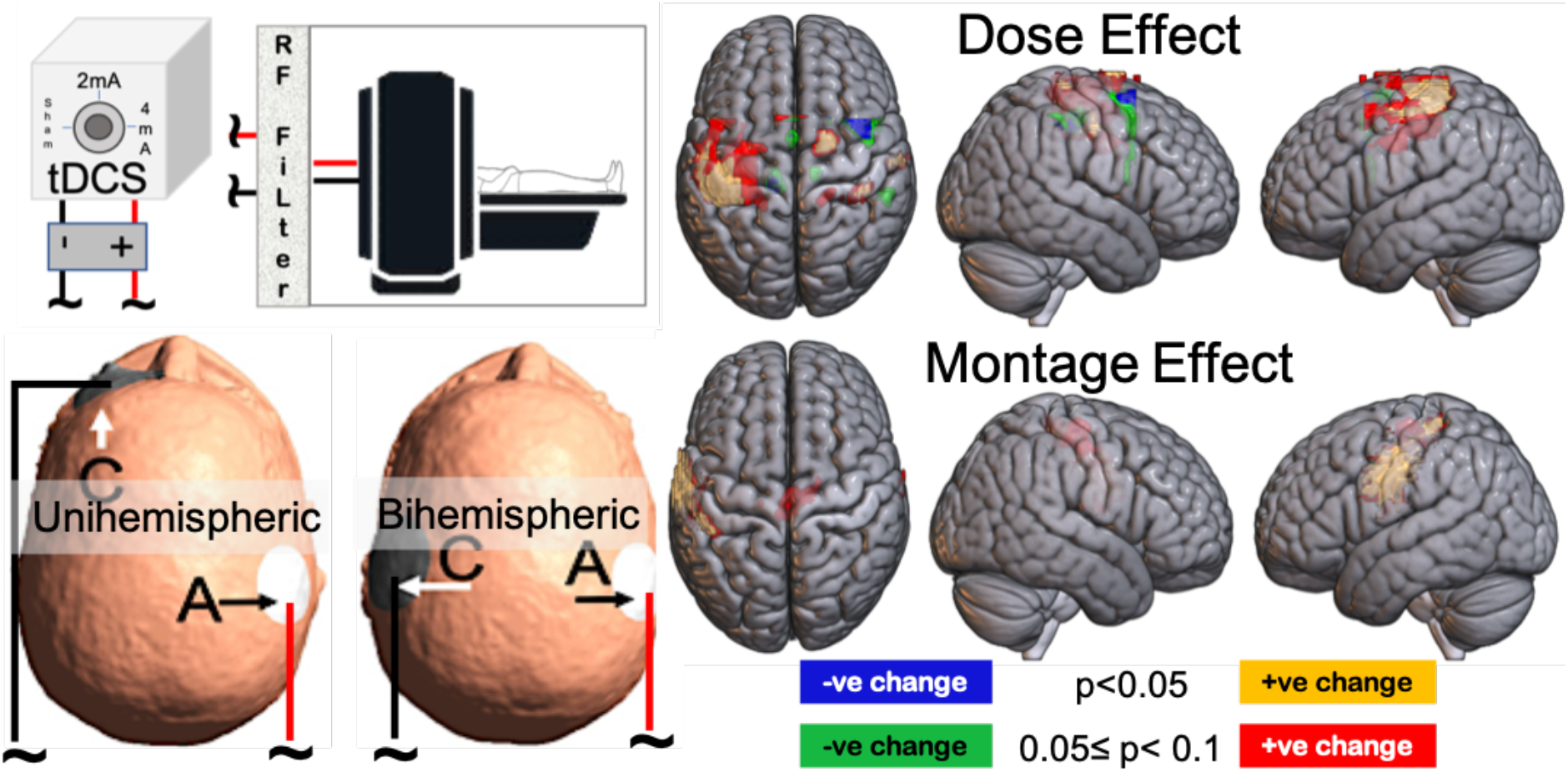

## 1. Introduction

Experimental studies have found evidence that tDCS modulates brain activity and affects behavior that draws on targeted brain regions (Stagg and Nitsche, 2011; Vines et al., 2006; Zheng et al., 2011). Prolonged sensory, motor, and cognitive effects of tDCS that outlast the stimulation period have been attributed to a persistent bidirectional modification of post-synaptic activity changes similar to long-term potentiation (LTP) and long-term depression (LTD)-like effects (Stagg and Nitsche, 2011). Due to these LTD- and LTP-like effects, tDCS or noninvasive brain stimulation has been thought of as a plasticity enhance of both directly- and remotely- targeted regions and of both functionally and structurally connected brain regions.

Depending on polarity, effects seen under the anodal or cathodal electrode differ. In a unihemispheric montage, the electrode of interest, either cathodal or anodal, is placed over the targeted area, while the reference electrode is typically placed over a region with minimal direct brain effect (e.g., supraorbital region on the contralateral side to the electrode of interest) (Bikson et al., 2009; Fregni et al., 2005; Nitsche and Paulus, 2000)(Bolognini et al., 2011; Lindenberg et al., 2010; Vines et al., 2008a). Furthermore, the return electrode is typically larger so that any residual effect onto any underlying brain tissue could be even further diminished. Contrary to these “unihemispheric montages,” a “bihemispheric montage” is typically chosen if the intent is to have both electrodes play an active role with the anodal electrode typically increasing the excitability of a targeted brain area while the cathodal electrode is thought to decrease the excitability of a region which might be a homotop region on the other hemisphere (Bolognini et al., 2011; Gomes-Osman and Field-Fote, 2013; Goodwill et al., 2016; Lindenberg et al., 2016; Waters et al., 2017). The bihemispheric montage studies (with up to 2mA current intensity) have been shown to be advantageous over unihemispheric montages in improving finger motor skills and facilitating motor learning in healthy subjects (Vines et al., 2008a; Waters et al., 2017) and in facilitating the motor recovery of stroke patients (Bolognini et al., 2011; Chhatbar et al., 2016).

Esmaeilpour and colleagues found incomplete evidence of a simple dose response for tDCS efficacy (Esmaeilpour et al., 2018). They highlighted that the tDCS dose effects studies had focused their attention on the current intensity range from 0mA to 2mA but saw a need to examine effects at higher doses to improve our understanding of the dose-response relationship (Batsikadze et al., 2013; Clark et al., 2012; Ho et al., 2016; Jamil et al., 2017; Zheng et al., 2011). Some studies over the last several years have expanded tDCS current intensity range up to 3mA (Agboada et al., 2020, 2019; Jamil et al., 2020) and even 4mA (Chhatbar et al., 2017), although more studies, in particular concurrent tDCS-fMRI studies, are necessary to examine relationships between stimulation dose and physiological signals (Esmaeilpour et al., 2020; Ghobadi-Azbari et al., 2020).

Some research studies investigated the safety, tolerability, and efficacy of 4mA dose in healthy adults and patients (Chhatbar et al., 2017; Dagan et al., 2018; Khadka et al., 2017; Trapp et al., 2019). Chhatbar and colleagues carried out a phase I dose escalation study showing safety and tolerability in stroke patients(Chhatbar et al., 2017). Khadka and colleagues showed that the high-intensity adaptive tDCS was tolerable in healthy subjects (Khadka et al., 2017). Trapp and colleagues demonstrated 20 sessions of adaptive 4mA tDCS for treatment-resistant depression (Trapp et al., 2019); Dagan and colleagues investigated the effects of multitarget tDCS with cumulative dose of 4mA on freezing of gait in patients suffering from Parkinson’s disease (Dagan et al., 2018). Additionally, Bikson and colleagues combined a current threshold value (0.63 mA/cm^2^) that caused brain damage in rats (Liebetanz et al., 2009) with a rat-to-human scaling factor to predict at what comparable dose level brain damage in humans could occur (Bikson et al., 2016). Based on this modeling, the threshold at which one could observe brain damage would vary from 67 mA to 173mA depending on the scaling factor used (Bikson et al., 2016). Our studies’ current is an order of magnitude lower than the current estimated by Bikson and colleagues to cause brain damage in humans.

In the current study, we examined whether: (1) regional cerebral blood flow (rCBF) in the targeted perirolandic region shows an effect of tDCS-DOSE applying up to 4mA with a total Charge Density (tCD) up to 0.18C/cm^2^ and current density of 0.31 mA/cm^2^; (2) electrode montage (unihemispheric versus bihemispheric) has a differential effect on rCBF; (3) a motor behavior such as finger sequence performance, drawing on the targeted/stimulated brain region, shows dose and montage effects; and (4) finite element modeling of current magnitude and flow correlates with behavioral and rCBF changes across tDCS dose levels.

## 2. Participants and methods

### 2.1. Participants

Thirty-two (32) healthy subjects (15 males, 17 females, mean age = 34.2 (SD =13.5)) participated in our single-blind study. One of the 32 participants was excluded from the analysis because of developmental brain abnormalities detected when MR images were examined. All remaining 31 participants were right-handed as assessed by the Edinburgh Handedness Inventory (Oldfield, 1971) and had no history of neurological or psychiatric conditions. The Institutional Review Board of Beth Israel Deaconess Medical Center approved this study, and all subjects gave written informed consent. Power analyses performed based on pilot data (Zheng et al., 2011) had indicated that we would need at least 10 subjects in each of the cells (three dose levels and two montages) to find an effect size of 1 for both dose and montage with a power of 0.8 at a two-sided alpha level of 0.05. Considering that we asked subjects to come back for six MR imaging studies, six finger sequencing experiments, and accounting for dropout, we oversampled slightly the total number of subjects suggested by the power analysis.

All subjects were asked to participate in six concurrent tDCS-MR imaging sessions and six concurrent tDCS-behavioral sessions. However, due to dropouts, including people moving away from the area and shutdown of non-clinical research studies due to the COVID19 pandemic, not all subjects were able to participate in all visits; details of the subject participation in different experimental conditions are collated in Table 1. Since not all subjects could complete all six behavioral sessions or all six MR sessions, we used linear mixed effects models to analyze the slightly unbalanced design.

**Table 1:** Visit count for different stimulation conditions in concurrent tDCS-fMRI and behavioral study.

|  | MRI | Behavioral |
| --- | --- | --- |
| Unihemispheric 0.1mA | 13 | 25 |
| Unihemispheric 2mA | 25 | 24 |
| Unihemispheric 4mA | 24 | 25 |
| Bihemispheric 0.1mA | 18 | 22 |
| Bihemispheric 2mA | 22 | 23 |
| Bihemispheric 4mA | 23 | 22 |

tDCS sessions for behavioral testing and imaging were separated by at least 24hours and were randomized in order. A portable single channel tDCS stimulator (neuroConn) was used for the behavioral sessions, while an MR compatible multi-channel tDCS stimulator (neuroConn) was used for the concurrent tDCS-MR sessions.

### 2.2. Electrode placement and stimulation dose

A round anodal electrode (diameter of 4cm) was placed over C4 (right precentral gyrus) for both the unihemispheric and bihemispheric montages while a round cathodal electrode (diameter of 5cm) was placed over either the supra-orbital region (corresponding to Fp1 in the unihemispheric montage) or over C3 (left precentral gyrus) in the bihemispheric montage. Stimulation sites (C3, C4, and Fp1) were localized using the 10-20 EEG measurement system and thoroughly cleaned with alcohol wipes. The rubber electrode (2mm thickness) was lathered with a 2-3 mm thick Ten20 conductive paste and pressed onto the scalp moving as much hair away under the electrode as possible to create the best conductive contact. Electrodes placed over C4 or C3 were held in place using a self-adhesive bandage and hypoallergenic medical tape when the electrode was placed at Fp1. Electrode connectors were adjusted to avoid any crossovers between wires. **Figure 1a** shows a representative electrode placement for both electrode montages.

**Figure 1:**
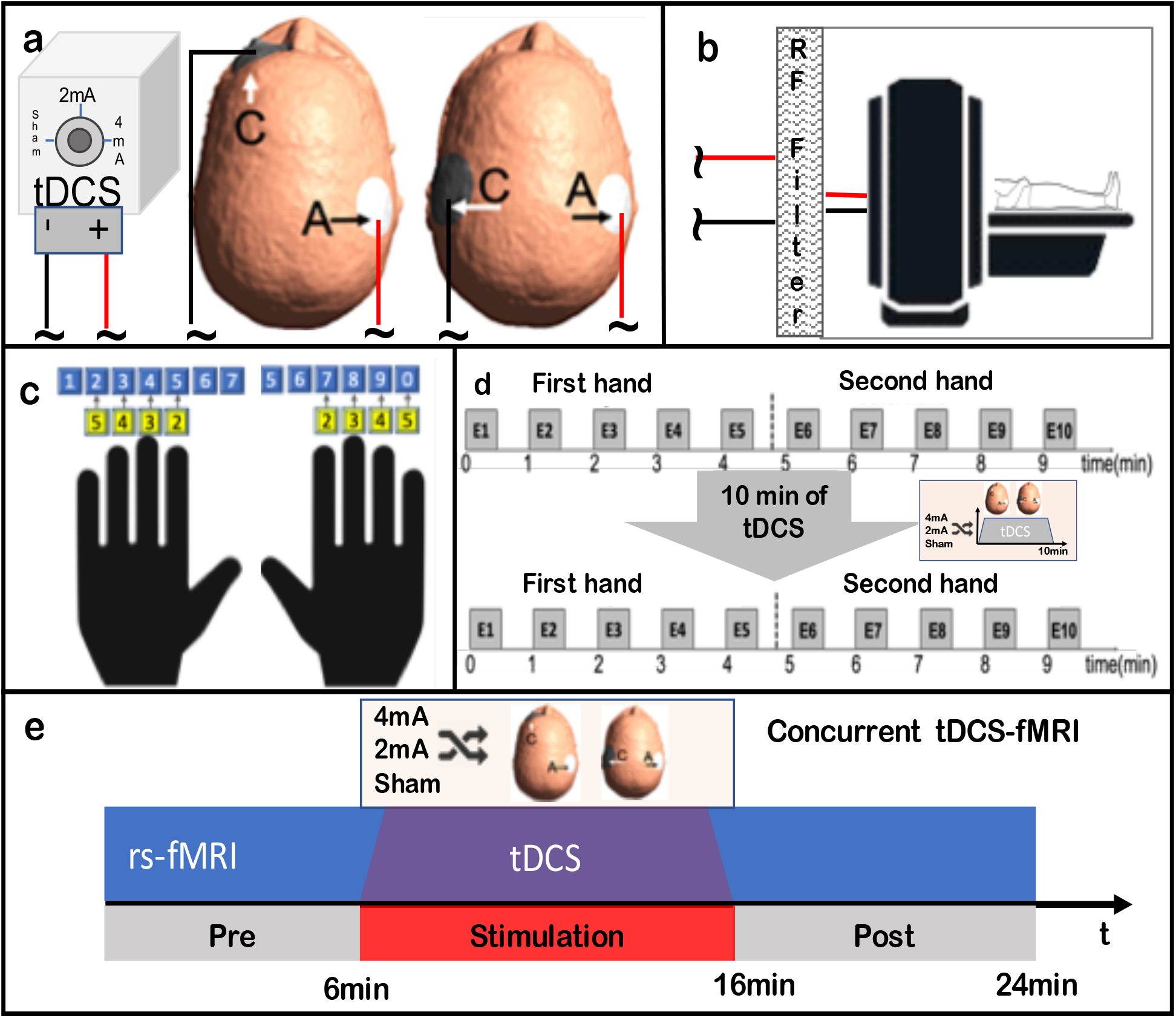
Schematics of experimental setup. (a)Head models show the two electrode montages used in this study and stimulator shows different dose levels. (b) Experimental setup for concurrent tDCS-fMRI using the RF filter panel to transmit the current into the MR room. (c) Finger-Key Mappings for left and right hand. Top row shows original keyboard keys with number pads pasted on the those keys displayed in the second row. (d) Behavioral task timing diagram: Finger sequence task performed before and after tDCS, the sessions consisted of 5 epochs (E=Epoch of 30seconds) for each hand before and after the tDCS stimulation; the hand that performed the finger sequence task first was counterbalanced across participants. (e) Concurrent tDCS-fMRI timing diagram: 24 minutes of continuous rs-FMRI was recorded during which 10 minutes of tDCS was applied from minute 6 to minute 16. The rs-fMRI timing diagram shown in 1e was the same for the ASL and the BOLD experiment which were done in one session, but the order of the ASL or BOLD experiment was counterbalanced as to which one occurred as first or second experiment; to simplify figure 1e, we only show the design for the ASL part of the combined experiment, each of them were 24 minutes long.

With regard to testing the effects of dose, our main aim was to compare 4mA (tCD = 0.1814C/cm^2^) with 2mA (tCD = 0.0910C/cm^2^) when applied for 10 min. We applied two 10-minute stimulations during MRI sessions one during ASL and the other during BOLD acquisitions, therefore tCD for those sessions were twice of that of behavioral study visits despite no change in the applied current intensity. We also developed a new sham-control condition, or quasi-sham condition, using a very low continuous direct current of 0.1 mA. This provided an initial sensory experience similar to other tDCS conditions while delivering a continuous current flow at a much lower tCD of 0.0045 C/cm^2^, which is less than 5% of the tCD of the 2mA condition. Several studies have used a low continuous current as a sham condition and reported that it is better at blinding participants to varying stimulation conditions than conventional ramping up and down sham conditions (Gibson et al., 2020; Trumbo et al., 2016). tDCS sessions were separated by at least 24hrs or more to provide for a wash-out period; physiological effects of tDCS have been reported to be no longer than a day for a single stimulation session (Agboada et al., 2019; Jamil et al., 2020).

### 2.3. Procedures

#### 2.3.1. Behavioral study

Each session started by placing electrodes onto the respective scalp locations as described above while the subject was sitting comfortably in a chair facing a computer screen and a standard keyboard placed on a table in front of the subject. The behavioral study involved a finger sequence task of each hand, performed as quickly and as accurately as possible over several epochs before and after a 10 minutes tDCS session (see Figure 1d). The finger sequence task was programmed using the software *Presentation* (Neurobehavioral systems, Berkeley, CA). A particular 7-digit finger sequence was randomly generated for each session. This first randomly generated 7-digit sequence was used to make each subject familiar with the task requirements in two 30 second periods using left and right hand. All subjects reported that two trials for either hand was enough to familiarize them with the task. After these warm-up trials, a new 7-digit sequence was generated for the actual task and this sequence remained the same for pre- and post-stimulation as well as for left and right hands. Subjects were instructed to type the 7-digit sequence repetitively as quickly and as accurately as possible in epochs of 30 seconds, separated by 30 seconds of rest and repeated five times before and immediately after 10 minutes of tDCS stimulation (see Figure 1d). The ordering for the hand that first performed the unimanual finger sequence task (left and right) was counterbalanced across participants.

##### 2.3.1.1. Task

Subjects were asked to use the index, middle, ring, and little fingers of the left hand (fingers 2-5) on keys 5, 4, 3, 2 and the corresponding fingers of the right hand on keys 7, 8, 9,10. The regular keyboard keys were taped over with numbers 2, 3, 4, and 5 for either the left or right hand (see Figure 1c) such that the index finger of both hands would correspond to number 2, middle finger to number 3, and so on. The task instructions for a single trial were to use the numbered keys from ‘2’ to ‘5’ on a standard computer keyboard to repeat a unimanual pattern of seven digits as quickly and as accurately as possible within 30-second epochs. The seven-digit sequence was randomly generated for each session and remained the same for pre- and post-stimulation.

During the task, the 7-digit sequence was displayed on a computer screen placed in front of the participant without any feedback. The experimental task for both hands lasted about 9 minutes before tDCS and also 9 minutes after tDCS. Each day of the experiment began with a short warm-up described above, which lasted about 3 min. All the keypresses and corresponding keypress times were recorded and exported to comma-separated values (CSV) files using the Presentation software.

#### 2.3.2. Concurrent tDCS-MRI methods

Prior to the MR procedures, all subjects were screened for contraindications to MRI. Subjects could not have any implanted electric, metallic, or magnetic material, and loose jewelry or other metallic objects were removed. The skin was inspected before and after electrode placement.

The device used for the tDCS-fMRI imaging study was an MR compatible direct current multi-channel stimulator (DCMC MR stimulator, neuroConn, NeuroCare Group, Germany), designed to perform noninvasive electrical stimulation inside the MR scanner. The DCMC MR stimulator was used to generate the electrical stimulation signal outside the MR scanner room. A computer program was used to generate the stimulation sequence used for concurrent tDCS-MRI. This sequence was started in parallel to the fMRI scan such that tDCS will be applied from minute 6 to minute 16 of the scanning (see Figure 1e). The rs-fMRI timing diagram shown in 1e was the same for the ASL and the BOLD experiment which were done in one session, but the order of the ASL or BOLD experiment was counterbalanced as to which one occurred as the first or second experiment; to simplify figure 1e, we only show the design for the ASL part of the combined experiment, each of them were 24 minutes long.

The stimulator signal is transferred inside the MR scanner room via a CAT5-DB9 9pin adapter on both sides of the RF panel (see Figure 1b). This is important, because any radio frequency (RF) noise from outside the MR scanner room could induce unwanted artifacts into the MR images. The adapter box in the control room (outside the MR scanner room) converts the multi-channel stimulation signal to a box cable compatible signal. On the other side, inside the MR scanner room the adapter box converts signal coming from an MR safe box cable into a stimulation signal which can then be applied to desired locations on a subject’s scalp using rubber electrodes. The last segment of the cable contained built-in 5 kOhm resistors as a safety precaution against cable resonances.

##### 2.3.2.1. MR Image acquisition

Each study visit involved an MR safety screening before the subject entered the MRI suite. Subjects were positioned in a Discovery MR750 3 Tesla MR scanner (GE Healthcare,Waukesha WI) in a supine position and images were acquired using a 32 channel RF head coil. Ear plugs and foam pads were used to minimize the noise and to help reduce head movements. Electrodes were positioned on the subject’s scalp prior to them entering the MR suite and the electrodes were connected with the DC stimulator cables once the subject’s head was positioned in the head coil. Calibration and localizer images were first acquired to assess the head orientation in the head coil. This was followed by acquisition of a T1-weighted (T1w) Magnetization Prepared fast gradient echo sequence with a 2mm slice thickness. This short T1w was used to check the electrode placement with reference to a targeted region of interest (ROI); electrodes were adjusted if necessary. The short T1w was repeated if needed before continuing with the remaining protocol. This was then followed by two separate 24 minutes long rs-fMRI scans, namely Arterial-Spin-Labeling (ASL) and Blood Oxygen Level-Dependent (BOLD)-contrast images. The BOLD sequence was a gradient-echo echoplanar imaging sequence with a TR of 3196ms, a TE of 24.0ms, a flip angle of 90, a FOV of 24cm, and a 2.5mm cubic voxel size. A total of 450 whole brain axial volumes were acquired. The 24 minutes long ASL resting state scan was recorded with a 3D volume difference image (tagged minus untagged) using a background-suppressed unbalanced pCASL scheme every 9 sec. Image resolution was 3.8×3.8 mm in plane and images had a slice thickness of 4mm with a FOV of 22cm. Labeling was performed 1 cm below the base of the cerebellum.

Separate ASL- and BOLD- RS-fMRI scans of 24 minutes duration each were recorded with 10 minutes of concurrent stimulation during each experiment as shown in the diagram (Figure 1e) for the ASL part of the experiment. The ASL and BOLD rs-fMRI order was counterbalanced to avoid order effects. Subjects kept their eyes open during both rs-fMRI scans, fixating on a green light on a screen visible through the head-coil mounted mirror and they were instructed to stay awake and let their mind wander, but not think about anything specific.

Transcranial direct current stimulation was applied for 10 minutes while acquiring resting state fMRI scans for 24 minutes, from minute 6 to minute 16 of each experiment, ramping up applied current from zero to the selected current level (sham 0.1mA, 2mA, or 4mA) in first 30 sec, and applied current levels were maintained for the next nine minutes, followed by 30 sec ramping down of applied current from the selected current level to zero. The diagram detailing the concurrent tDCS rs-fMRI protocol for one part of both experiments is shown in Figure 1e.

Prior to the start of our study, we tested the induced noise and artifacts on phantoms and eventually a volunteer. We found, unsurprisingly, that RF filtering of the wires from the current controller was essential to avoiding elevation of noise with concurrent tDCS-MRI. There was no sign of increased noise or a change in signal to noise ratios when the current was on compared to off or when higher currents (e.g., 4 mA) were applied compared to 2mA or sham.

Further, a magnetization prepared fast gradient echo sequence (a high resolution T1 weighted, BRAVO sequence from GE) with a TR of 2400.0 ms, a TE of 3.36ms, FOV 24cm, a 1×1×1mm^3^ voxel resolution, flip angle of 8, fat suppression, and a TI of 1000ms was acquired. A 3D high-resolution T2w image was acquired as well with a TR of 5000.0ms, a TE of 96.0ms, a FOV 24cm, and a voxel resolution 1×1×1mm^3^. These high-resolution scans were acquired in only one of the MR sessions after removing the electrodes in order to get the best estimate of the scalp thickness. The high-resolution T1w and T2w images were acquired only once per subject, and they were used for modeling the electric field.

#### 2.3.3. Safety and tolerability

After each session of tDCS, we recorded safety and tolerability information. The skin and scalp location under the electrodes were inspected for any skin burns or other lesions. At the end of each session, we asked volunteers to indicate their tolerance of the noninvasive brain stimulation on a visual analog scale (VAS) with 0 and 10 as the endpoints where zero indicated that subjects tolerated the stimulation well and had no unusual sensations and 10 indicated that the stimulation session caused strong sensory experiences and was judged to be barely tolerable. At no point were subjects provided any feedback on whether or not they had received high dose, lower dose tDCS or sham tDCS.

### 2.4. Data Analysis

#### 2.4.1. Behavioral study

A custom MATLAB^®^ (Mathworks, Natick, MA, USA) script was used to analyze the finger sequence task’s performance measures. The keypresses and keypress times were used to extract the number of correct and partially correct 7-digit sequences. A partially correct sequence is a sequence that has at least 4 or more consecutive keys of the 7-digit sequence correctly entered and gives partial credit accordingly (e.g., 4/7, 5/7, 6/7). In addition to this accuracy measure, we also calculated inter-keypress times, the standard deviation, and the variance in inter-keypress times.

Hashemirad and colleagues, in their review of tDCS effects on finger sequence learning, suggested that besides changes in movement speed and accuracy, a skill measure, a combination of speed and accuracy, could be considered as behavioral outcome measures to monitor improvement in tDCS-induced motor performance changes (Hashemirad et al., 2016). Thus, we calculated a measure of both accuracy and time by dividing the sequence count by the standard deviation (SD) of the inter keypress time. The sequence-count is the sum of partially correct and correct sequences (sequence count) in each epoch and the SD of keypress time is the standard deviation of inter keypress times during each epoch. We also calculated an average change in the “sequence-count/SD of keypress time” over five epochs (see Figure 1d) comparing before and after tDCS. For this study, we refer to the change in “sequence-count / SD of keypress time” as a change in FingerSequencePerformance (FSP).

A preliminary analysis of the entire dataset showed that there were epochs across subjects and conditions that had an excessive number of errors. To avoid skewing the analysis with these outlying data, we developed a very conservative regimen for eliminating outliers. Outliers were identified as 30s epochs for which the number of errors was greater than two SDs above the mean number of errors across all 30s epoch periods. If an epoch was identified as an outlier, we removed that particular epoch from further analyses. Out of 2820 epochs, we identified 41 outliers in total (less than 1.5% of all epochs), including 24 pre-stimulation outliers. FSP is expected to improve from before stimulation to after stimulation in each visit due to practice effects. However, we observed that some participants’ change in FSP was lower than their pre-stimulation FSP. This was due to different reasons such as decreased attentiveness, misreading the instructions, and failure of the presentation software or the computer. We discarded 11 data points that showed a negative change in FSP of more than 100%.

A preliminary descriptive analysis of the FSP data revealed that the change in FSP was affected by the session order when data were collapsed across dose and montage; change in pre to post stimulation “seqcnt_by_sdtime” dropped continuously from visit number 1 to 4, and it either stabilized or improved again slightly for visit number 5 and 6. Because these effects changed with the visit number, this can be attributed to practice effects. Reports in the literature suggest that behavioral and physiological effects of short tDCS stimulation last no longer than a day (Agboada et al., 2019; Jamil et al., 2020); since the task was performed before and immediately after stimulation on every visit, which was separated by 24hrs or more, visit-wise changes in the dependent variable “seqcnt_by_sdtime” for the task performed before stimulation is representative of practice effects. In order to isolate practice effects across visits, we calculated an average pre-stimulation FSP value for each visit, from visit number one to six, and normalized these values by dividing them with the maximum found in any of the 6 visits. To get rid of practice effects we divided each pre-post FSP change by the associated normalized practice effect based on the visit number.

#### 2.4.2. Concurrent tDCS-MRI study

Although we recorded BOLD and ASL rs-fMRI images, here, we only report the ASL data. BOLD data were mainly acquired to examine functional connectivity, which will be reported elsewhere.

##### MR Image Preprocessing

For each session, the low resolution T1w structural scan was first segmented into gray matter, white matter, cerebrospinal fluid (CSF), and coregistered to MNI space with the Computational Anatomy Toolbox (CAT12). The 4D ASL images were preprocessed with FSL and SPM12. First, 4D ASL images were split timewise into 160 individual NIfTI volumes using fslsplit. NIfTI volumes were realigned to the mean image using SPM12 and newly realigned images were written out, using a quality of estimation at 0.9, sampling distances in reference image at 4mm, and using 7th degree B-spline interpolation for estimation and reslicing. Mean image and realigned images were coregistered to the subject specific gray matter volume (created using CAT12 toolbox). The transformation matrix generated by the CAT12 toolbox while coregistering the T1w image to MNI space was applied to normalize all the coregistered rs-fMRI images to MNI space. Normalized images were smoothed using an 8mm FWHM isotropic Gaussian smoothing kernel.

##### Image Analysis

At the first level, perfusion ASL images were fitted into an ON- vs. OFF-stimulation [1 −1] model examining effects of 2mA and 4mA dose with unihemispheric and bihemispheric montages. The first minute of each ASL acquisition was excluded from the analysis to allow for stabilization of the signal. We calculated a T-contrast image comparing 10 minutes of an ON period with the OFF periods preceding and following the ON period (15 min). Stimulation condition effects were estimated using a second-level general linear model (GLM) in SPM12 by using first level contrast images as inputs across all subjects who participated in that stimulation session. Global differences in scan intensity were removed by scaling each scan in proportion to its global intensity. Since the number of participants in the stimulation conditions were not the same and in order to have uniformity across stimulation conditions, we transformed group level T contrast images to Fisher transformed z-images using a custom MATLAB script.

##### ROI definition and analysis

In a 2nd level analysis across all subjects and all stimulation conditions (unihemispheric and bihemispheric montages with the 2mA and 4mA dose), a region with suprathreshold activation was identified in the right perirolandic region (Figure 3) that was common across all stimulation conditions. A 3D ROI was created by including those voxels that survived a strict threshold (p_unc_<0.01). We mirrored this 3D ROI from the right hemisphere to the left hemisphere to give us two ROIs of equal size centered on the perirolandic regions on either hemisphere. These two ROIs were then used to extract ROI-averaged T-contrast values specific to each stimulation session’s first level whole brain SPM analysis.

##### Linear mixed effects analysis

For each dependent variable (FSP, left ROI ASL T-Values, and right ROI ASL T-Values), a linear mixed effects model was run with two fixed effects, namely montage and dose, and one random effect, which was subject. A mixed effects model is appropriate here as multiple conditions were run for each subject, but not every subject participated in all conditions, thus using subject as random effect control for intersubject variability and missing subject-condition values. R was used for all mixed effects model calculations. The package *lme4* was utilized for mixed effects calculation, and the package *lmerTest* was utilized to calculate F scores and p values. The *lmerTest* package uses the Satterthwaite degrees of freedom calculation method, which can then provide the associated F scores and p values, which are reported here. For all mixed effects models that found a main effect of dose, post-hoc corrected multiple comparison tests were also run using estimated marginal means to see which dose levels differed from which. The R package *emmeans* was used for all post-hoc tests. A Tukey correction was used for all multiple comparisons, and the Kenward-Roger method for calculating degrees of freedom, and thus t values and p values, was used.

#### 2.4.3. Tolerability analysis

The tolerability data recorded from both Behavioral and concurrent tDCS-MRI sessions was analyzed separately using linear mixed effects models. Tolerability scores for behavioral sessions were correlated with the left and right FSP. Similarly, concurrent tDCS-MRI tolerability scores were correlated with ASL T contrast values for each session.

### 2.5. Electric Field Modeling

It has recently become possible to simulate the electric field distribution through the brain with finite element modeling and there are several software routines available for this modeling approach (Huang et al., 2019; Laakso et al., 2016; Saturnino et al., 2017). To prepare our data for the modeling, we used an application that is available through a freely available software package called SimNIBS 2.1 (Saturnino et al., 2017). We performed a skull segmentation on the high resolution T1w and T2w images for each participant using the *unified segmentation* routine implemented in SimNIBS, which combines spatial normalization to MNI space, intensity inhomogeneity correction, and tissue segmentation into one model. The segmentation routine uses a Gaussian mixture model for modeling tissues using the spatial tissue probability map from SPM12. The distribution of the electric field potential (V/m) is influenced by the placement of the electrodes, the inter-electrode distance, and the current dose. To mimic the exact electrical field applied during the MR imaging experiment, we identified electrode locations from the short T1w image acquired before recording the ASL rs-fMRI images. These electrode locations were then transferred to the native space of the high resolution T1w image. We generated electric field models representing the magnitude of the electric field potential at each voxel in the simulated brain model based on electrode locations, stimulation dose-montage information, and segmentation output. A custom MATLAB script was used to generate a brain volume with each voxel representing the electric field (V/m) and transform them into MNI space. Group level stimulation effects were calculated by averaging the simulated brain volumes. The ROI from the ASL analysis (see 2.4.2) was used to extract voxel-wise values from the group level electric field model.

### 2.6. Correlation analyses

Scatterplots and correlation analyses were used to investigate correlation between the observations. There are four groups of correlations that were analyzed. 1) ASL T Contrast from left- and right-ROI against FSP values from left- and right- hand. 2) Tolerability scores in MR experiments against ASL T-contrast and Tolerability scores in Behavioral experiment against FSP values. 3) Left- and right- FSP values against simulated current intensity values from left- and right perirolandic ROI. 4) Left- and right-ASL T-Contrast values against simulated current intensity values from left- and right perirolandic ROI. For each group, there were four pairs of correlations calculated. Correlation values and significance were calculated for each of these parameter pairs using the Pearson’s r method.

## 3. Results

### 3.1. Dose and Montage Effects: Finger Sequence Performance (FSP) and regional Cerebral Blood Flow (rCBF)

Figure 2 shows the unilateral FSP for each stimulation condition. LME analysis with DOSE and MONTAGE as main factors showed a significant effect of the factor DOSE for both right hand FSP(F(2,97)=9.28, p=0.0002) and left hand FSP(F(2,96)=6.30, p=0.0026). Factor MONTAGE and the MONTAGE-DOSE interaction did not show any significant effect on FSP for either hand. For the significant main effect of dose on left-hand FSP, sham differed from both 2 mA and 4 mA (t(101)=−2.678, p<0.05, and t(102)=−3.230, p<0.005, respectively). For the significant main effect of dose on right-hand FSP, sham also differed from both 2mA and 4mA (t(101)=−2.628, p<0.05, and t(103)=−4.155, p<0.0005, respectively).

**Figure 2:**
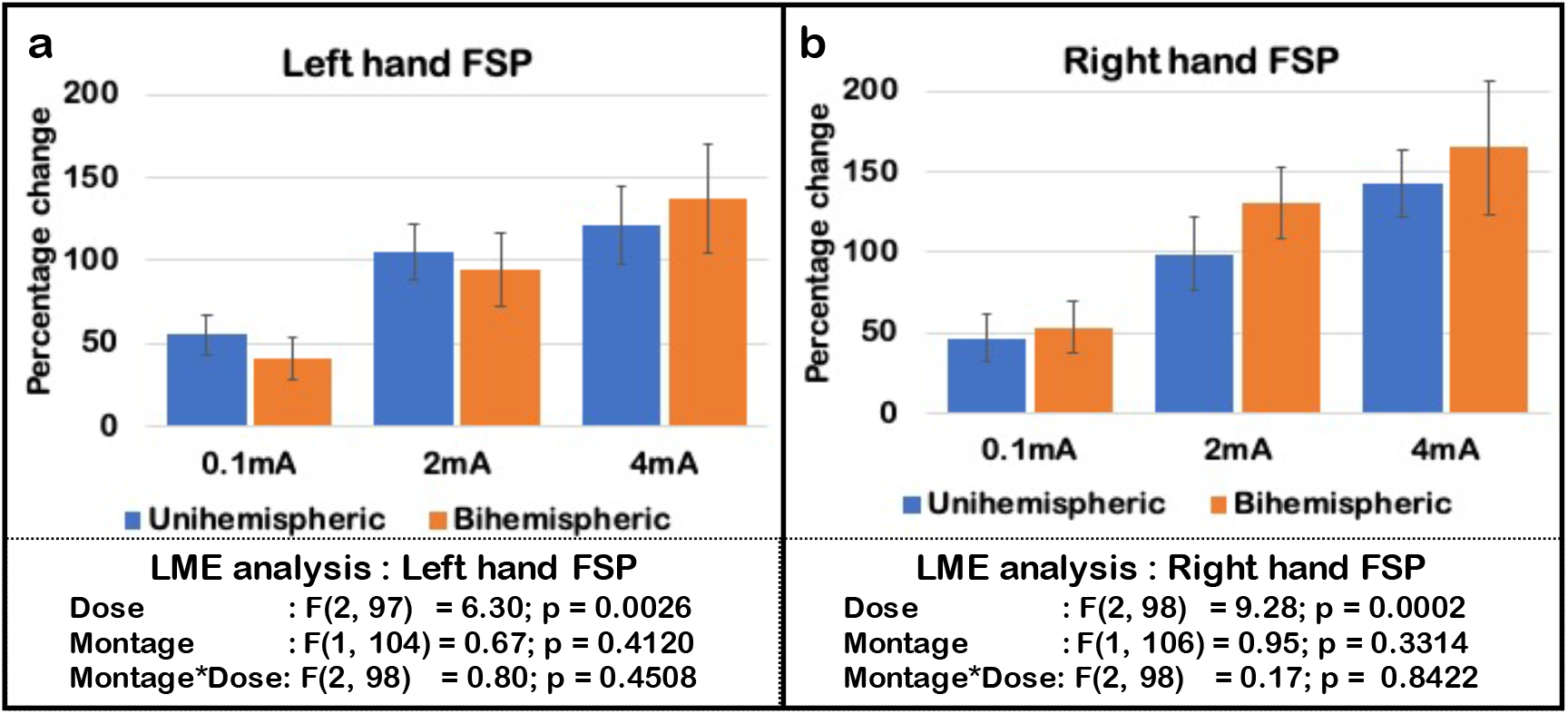
Percentage change in Finger Sequence Performance (FSP) comparing before to after stimulation (MEAN +/− SEM). (a)Left hand FSP change and (b)Right hand FSP change.

**Figure 3:**
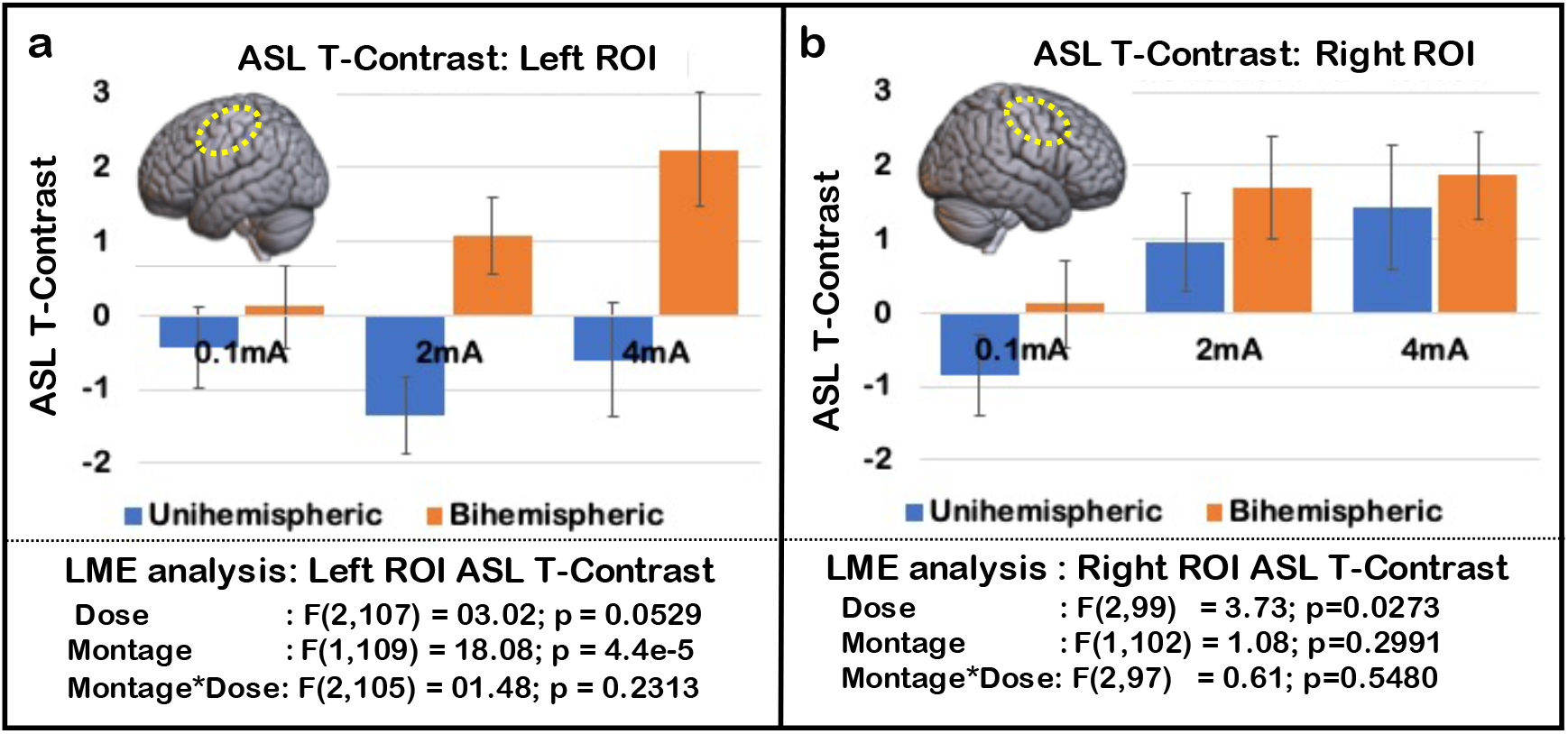
ASL T contrast values in Left (3a) and Right (3b) perirolandic ROI (MEAN +/− SEM) comparing ON versus OFF periods of tDCS stimulation.

Figure 3 shows the rCBF changes recorded as ASL T-Contrast values. LME analysis showed a significant effect of dose on the right ROI ASL T-Contrast values(F(2,99) = 3.73, p=0.0273) and significant effect of montage on the left ROI ASL T-Contrast values (F(1,109) = 18.08, p = 0.00004). The effect of dose on left ROI ASL T-Contrast showed a trend (F(2,107) = 3.02, p=0.052). For the significant main effect of dose on the right ASL contrast, sham differed from 4mA (t(107)=−2.583, p<0.05).

### 3.2. Safety and tolerability

A total of 266 tDCS sessions were administered. None of our participants had any significant adverse effects (e.g., severe headaches, seizures, neurological impairments, skin burns, or any hospitalizations directly related to the high dose tDCS stimulation), and we did not detect any MR artifacts due to applying high dose tDCS inside the scanner bore and the head coil underscoring the safety of the MR compatible system delivering transcranial direct current in the MR machine. Using the RF panel to transfer the current into the scanner room allowed us to filter out any RF from outside before entering the scanner room and also any RF from inside to escape outside. We were able to obtain MR images without any distortion, signal loss, or signs of elevated flip angles near the electrodes or due to the tDCS stimulation. The total impedance between two tES connectors connected to anodal and cathodal electrodes was continuously monitored.

Almost all subjects experienced an initial tingling, itching, and/or warming/burning sensation when the current was ramped up, which typically became less intense or subsided after this initial ramp up. None of the subjects indicated that any of the sessions were barely tolerable or intolerable to them. However, the data showed that current dose had a strong effect on the tolerability scores with 4mA stimulation conditions having higher scores than 2mA and sham 0.1mA (Figure 4) across the unihemispheric and bihemispheric montages. The bihemispheric sham condition stood out as having the lowest scores on the visual-analog tolerability scale. LME analysis showed a significant effect of dose on tolerability scores reported by subjects in both behavioral (F(2,108)=67.94, p<2e-16) and concurrent tDCS-MRI experiments(F(2,98)=38.65, p<4e-13). For the significant main effect of dose, tolerability with sham stimulation was significantly different from 2mA and 4mA in both experiments: (t(113)=−7.07, p<0.0001 and t(113)=−11.26, p<0.0001 respectively for the behavioral experiment, and t(105)=−6.01, p<0.0001 and t(107)=−8.44, p<0.0001 respectively for the MR-concurrent experiment). Tolerability scores differed between 2mA and 4mA stimulation conditions: (t(112)=−4.16, p=0.0002 for behavioral, and t(102)=−2.75, p=0.019 for concurrent).

**Figure 4:**
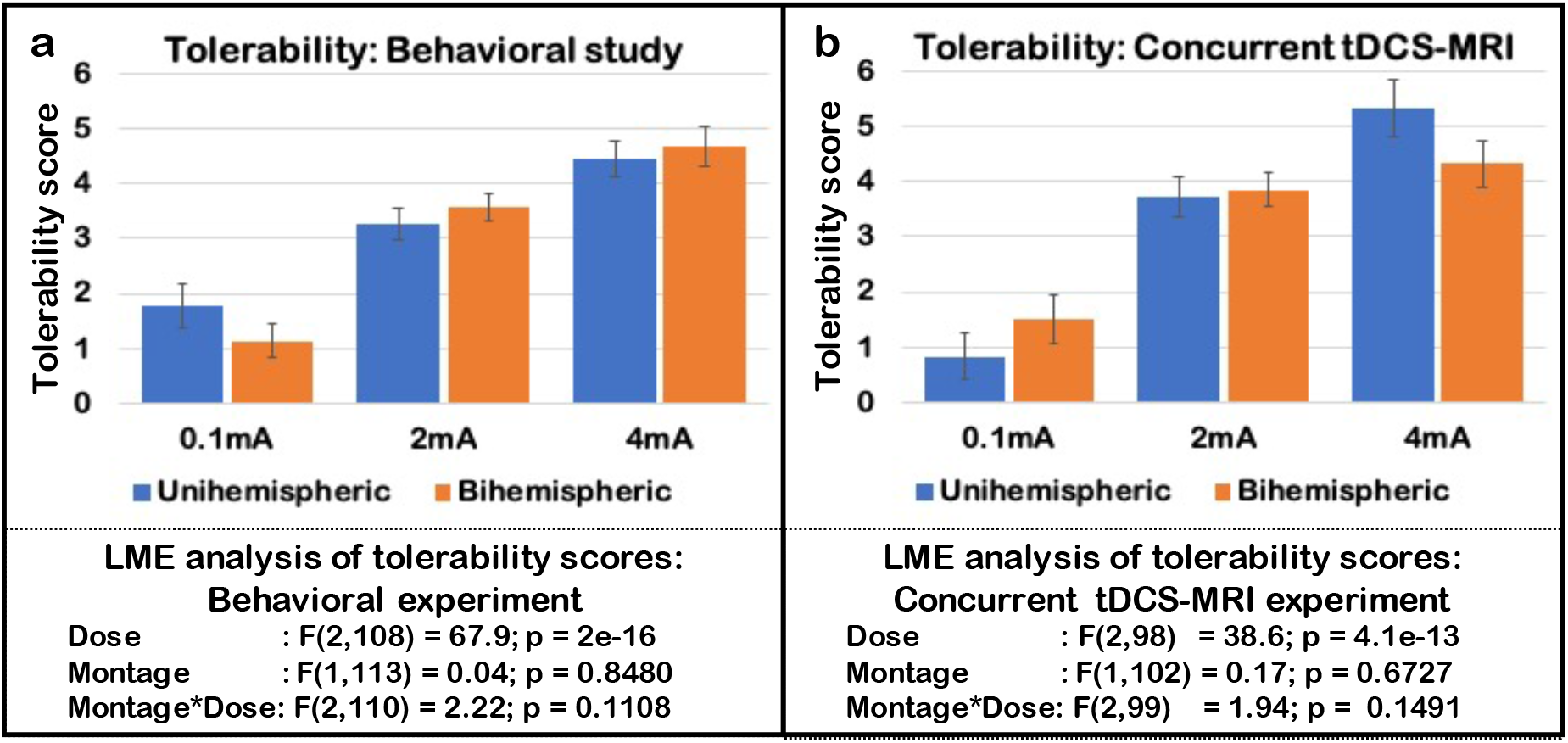
tDCS tolerability analysis in a) behavioral study and b) concurrent tDCS-MRI study.

### 3.3. Dose and Montage Effects on regional Cerebral Blood Flow: SPM-Maps

Figure 5 shows the second level analysis results for all active stimulation conditions, namely Unihemispheric 2mA, Unihemispheric 4mA, Bihemispheric 2mA, and Bihemispheric 4mA. For illustration purposes and comparison across conditions, we thresholded all group level analyses images at uncorrected p-value of 0.05 as well as p-value of 0.1, and restricted the analysis to a part of the brain that consisted of the pre- and post-central gyrus (i.e., perirolandic region), the adjacent premotor region, and the frontomesial region. The unihemispheric 2mA montage showed positive rCBF changes in the right perirolandic region and negative rCBF changes in the left perirolandic region; positive right perirolandic changes were slightly more spatially dispersed in the 2mA condition. Comparing 4mA to 2mA showed higher blood flow changes in perirolandic regions on both hemispheres and in the left premotor region. The bihemispheric montage produced positive rCBF changes in the right perirolandic region for the 2mA condition, and in an extensive bilateral pattern for the 4mA condition involving the perirolandic region (upper and lower parts) and frontomesial regions. The bihemispheric 2mA caused negative rCBF changes in the left perirolandic region (exposed to cathodal stimulation). In contrast, bihemispheric 4mA stimulation showed only small negative rCBF changes in the bilateral premotor area. Comparing bihemispheric 4mA to 2mA showed positive rCBF changes in the upper part of the bihemispheric perirolandic region.

**Figure 5:**
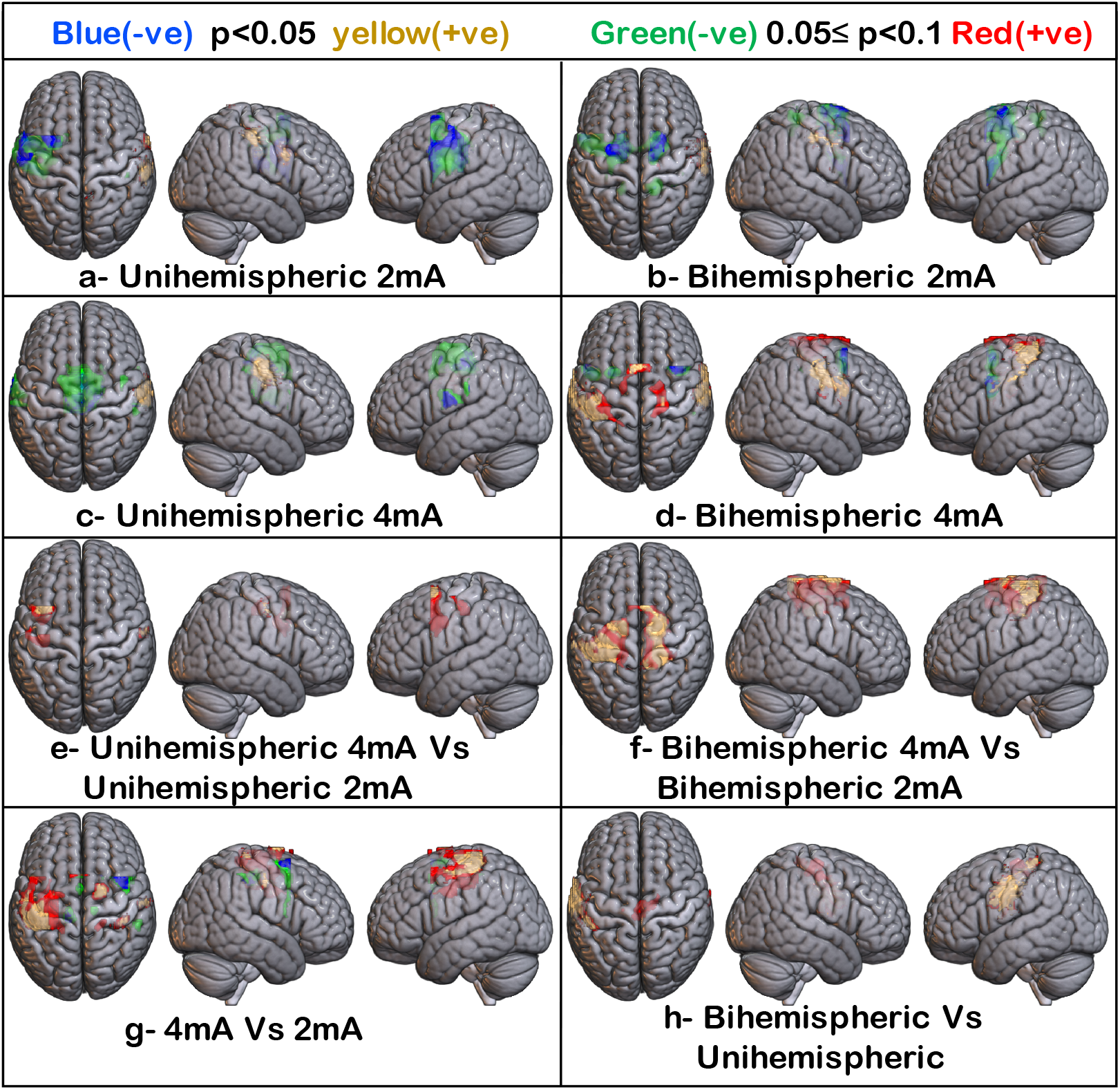
SPM Activation maps for all the active stimulation conditions with ON>OFF stimulation function and for dose-montage effects; (a, c, and e) Unihemispheric montage: SPM maps for 2mA, 4mA, and 4mA vs. 2mA dose. (b, d, and f) Bihemispheric montage: SPM maps for 2mA, 4mA, and 4mA vs. 2mA dose. g) Effect of Dose with function 4mA>2mA, and h) Effect of Montage with function Bihemispheric montage>Unihemispheric montage. Uncorrected threshold of p=0.05 was used to identify positive and negative changes in ASL represented by Yellow and Blue voxels, respectively. Red and Green voxels represent positive and negative changes in ASL values for p-values between 0.05 and 0.1).

Interactions between dose and montage using SPM were done for the 4 active stimulation conditions, namely unihemispheric 2mA, unihemispheric 4mA, bihemispheric 2mA, and bihemispheric 4mA (Figure 5). The 4mA dose (both montages) level when contrasted against 2mA showed positive rCBF changes in the right and left perirolandic region and in bilateral frontomesial regions, while it showed negative changes in the right perirolandic and right premotor region (Figure 5g). The positive rCBF effect of electrode montage, bihemispheric vs. unihemispheric, was mainly observed in the left perirolandic region. There were no negative changes in rCBF comparing the two montages.

### 3.4. Correlation between rCBF, Finger Sequence Performance, simulated current density, and tolerability scores

Across dose conditions, changes in both left and right hand FSP did not correlate with rCBF changes in the left or right perirolandic ROI (Figure 6a). Tolerability scores obtained after the concurrent tDCS-MRI experiments did not correlate with regional ASL T- Contrast values (Figure 6b). NormJ values of left and right ROI were correlated with four output variables (FSPLeft, FSPRight, ASLeft, and ASLRight); Bonferroni correction for multiple comparisons set an **α**=0.0125. Moreover, tolerability scores were correlated with two outcome variables each setting the multiple comparison corrected **α** value to 0.025. FSP Right values significantly correlated with the simulated current intensities(normJ) of both left and right perirolandic ROI (Figure 6c). ASL T- Contrast values from left and right ROI showed no significant correlation with normJ values (Figure 6d) with the best correlations seen between the right ROI ASL T-Contrast and normJ values (r=0.13; p=0.15).

**Figure 6:**
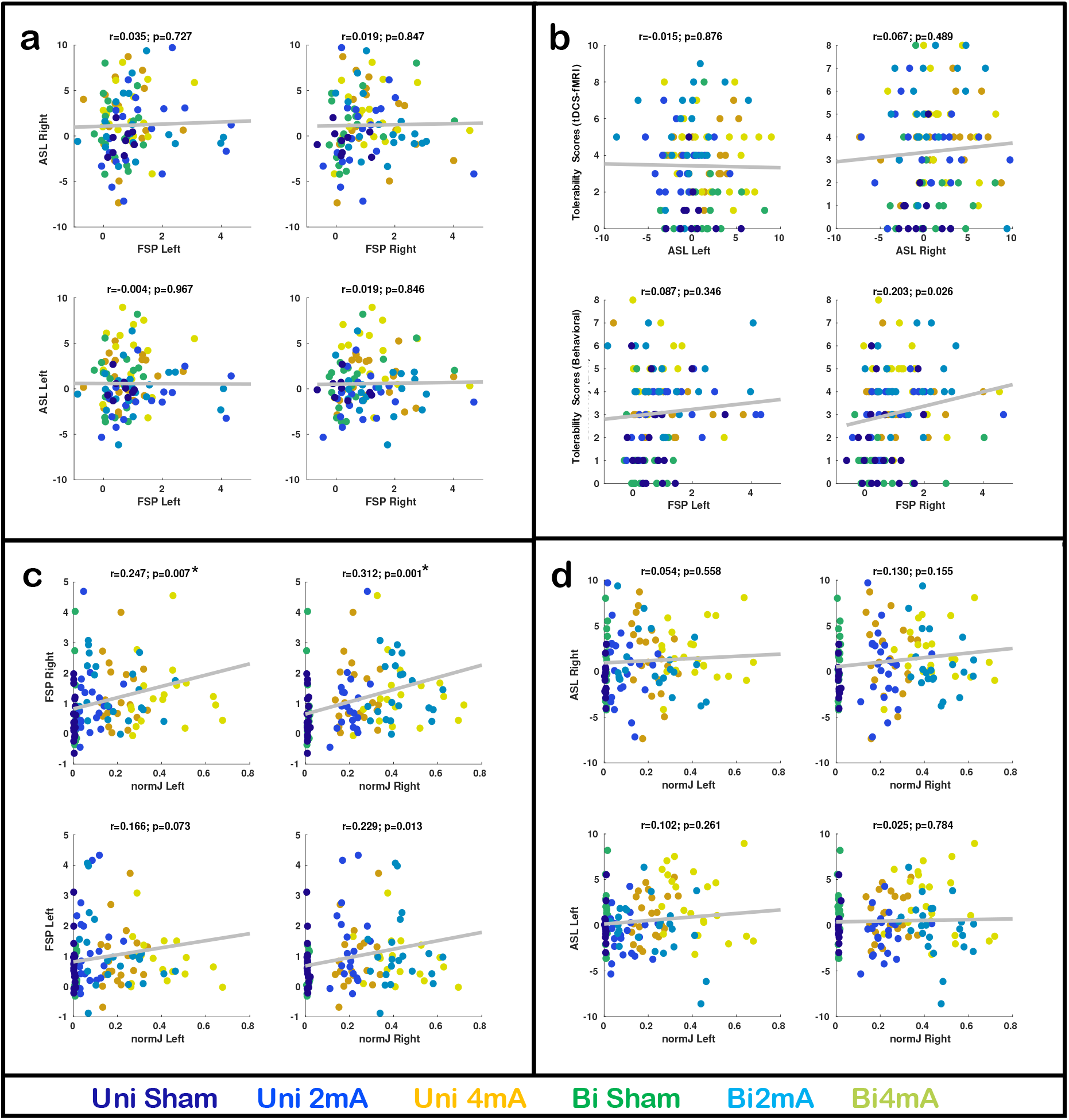
Correlations between observed output variables. a) left and right ROI ASL T-contrast values against FSP plotted across all dose-montage combinations. (b) Tolerability scores against ASL T-Contrast and FSP values. (c) FSP values from left and right hand against simulated current intensity from left and right perirolandic ROI. (d) left and right ROI ASL T-Contrast against simulated current intensities. *-multiple comparison corrected significant correlation.

## 4. Discussion

The most important findings of our studies were that the 1) finger sequence performance showed a linear tDCS-DOSE effect for both the left and the right hand but no effect of montage; 2) regional blood flow changes in the left perirolandic region showed significant effects of tDCS-montage and nearly significant effect of tDCS-dose while rCBF in the right perirolandic region (anodal electrode) showed a significant linear trend for tDCS-DOSE. In particular, the 4mA cathodal electrode over the left perirolandic region led to an increase in rCBF compared to 2mA bihemispheric montage. The two highest rCBF increases were seen with the 4mA cathodal electrode over the left perirolandic region and with the 4mA anodal electrodes over the right perirolandic regions, in the bihemispheric and unihemispheric montages respectively; 3) FSP changes of both hands correlated with simulated current density in the perirolandic ROI from contralateral hemisphere and FSP changes in right hand also correlated with simulated current density from right hemispheric perirolandic ROI; 4) strong right hand FSP changes in the condition with anodal electrode over the right perirolandic region suggest that ipsilateral motor pathways could be activated more at higher current doses; and 5) finger sequence performance of left hand, regional blood flow changes in the left- and right-perirolandic ROI did not correlate with the tDCS session tolerability scores.

### Finger Sequence Task

Finger sequencing tasks have been used to study effects of tDCS (Karok and Witney, 2013; Vines et al., 2008a, 2008b; Waters-Metenier et al., 2014). We used a longer sequence of 7 digits instead of the more commonly used shorter 5-digit sequence, since a 7-digit sequence with 4 keyboard keys allowed us to create 576 unique sequences compared to 96 unique sequences possible with 5-digit sequence. This significantly reduces the possibility of a participant being assigned similar sequences more than once in the randomization process. In addition, difficulty level increases with the length of a sequence, reducing the possibility of ceiling effects in the finger sequence task. Evidence suggests that studying either speed (inter-keypress times) or accuracy alone provide only a limited understanding of the tDCS stimulation effects (Hashemirad et al., 2016; Reis et al., 2009). Our analysis used accuracy reflected by the correct sequence count and a time variable reflected by the standard deviation of keypress times into one outcome variable called finger sequence performance (FSP). A review of the published tDCS-finger sequence task studies found that anodal tDCS showed improvement in accuracy compared to sham and no significant difference in accuracy between unihemispheric and bihemispheric tDCS (Hashemirad et al., 2016). In our study, we observed that increasing dose significantly impacts behavioral outcomes in both left and right hands. Both hands show improvement in the FSP with the increase in dose levels. The literature suggests that there is no significant difference in the sequence learning accuracy after application of either unihemispheric or bihemispheric montage. Our results confirm these earlier findings with a typical dose of 2mA and extend this finding up to the higher dose levels of 4mA.

### Dose and Montage Effects

Dose and montage of noninvasive brain stimulation have been used in the past to show that modulation of tDCS parameters can lead to differential effects in behavioral outcome measures (Bolognini et al., 2011; Hashemirad et al., 2016; Jamil et al., 2020; Vines et al., 2006). A dose of the applied/injected current has been of particular interest since there is concern that a certain amount of the current might be shunted to the return electrode through the scalp and might not enter the brain. This has prompted the search for surrogate brain markers that could be used to examine variations in tDCS parameters such as dose, montage, and duration. Regional CBF changes under the anodal and to a slightly lesser degree under the cathodal electrode have shown a characteristic on-off pattern and a significant correlation with the applied current levels, although current doses of less than 2mA have mostly been examined in the past (Antal et al., 2014, 2011; Lin et al., 2017; Meinzer et al., 2014; Stagg et al., 2013; Zheng et al., 2011). Some tDCS studies have already reported stronger effects with higher current doses without jeopardizing safety and worsening tolerability (Jamil et al., 2020; Liu et al., 2018; Trapp et al., 2019). tDCS-ASL studies (Jamil et al., 2020; Stagg et al., 2013) have reported increase in perfusion with the electrode montage similar to the unihemispheric stimulation montage used in our study. Our findings concur with these earlier studies that investigated effects of Anodal tDCS. These tDCS-ASL studies have also shown the decrease in perfusion with unihemispheric montage- cathodal stimulation; unfortunately we did not investigate such polarity dependent effects. Instead, we investigated both unihemispheric and bihemispheric electrode montages with anodal electrodes on the right perirolandic region lead that lead to an increase in rCBF with higher dose.

Several studies have now shown that the common assumption that the cathodal electrode in a bihemispheric montage might lead to a detriment in performance or a temporary cortical dysfunction might not be true in every situation (Ciechanski and Kirton, 2016; Vines et al., 2008b; Waters-Metenier et al., 2014; Waters et al., 2017). They also showed that the advantages of a bihemispheric montage over a unihemispheric montage persisted even when the polarity of the bihemispheric montage was reversed. Waters and colleagues (2017) postulated an ipsilateral control of motor actions and that bihemispheric tDCS (2 mA, 25 minutes) could lead to more cooperation between hemispheres and less transcallosal inhibition (Waters et al., 2017). Our concurrent tDCS-fMRI study provides some support for the notion that the cathodal electrode in a bihemispheric montage could act as an excitatory stimulation instead of inhibitory. This will need to be examined further in other experiments in the future. In another study, Batsikadze and colleagues observed increase in Motor Evoked Potential (MEP) values when 2mA cathodal tDCS stimulation was applied to left primary motor region for 20 minutes(Batsikadze et al., 2013). Although the aforementioned study used unihemipsheric electrode montage the results show that higher intensity cathodal tDCS may actually lead to exitatory effects, also suggested by our investigations into tDCS-induced rCBF changes (see also Zheng et al., 2011).

Montage design has also attracted more attention as a way of optimizing an electrode placement to engage or target particular brain regions and to model this regional engagement by using different multi-electrode montages that ensure that a particular brain region receives the desired current dose. Electrode montage and in particular, a bihemispheric montage with transcallosal interactions might have a more complex effect on brain function. Regional cerebral blood flow as a noninvasive physiological surrogate of brain and synaptic activity showed a clear positive dose effect underlying the anodal electrode in the unihemispheric and bihemispheric conditions, but only a prominent increase in rCBF underlying the cathodal electrode in the 4mA bihemispheric condition. A similar non-linear effect with higher tDCS dose was seen in frontomesial brain regions with the 4mA bihemispheric montage. The frontomesial regions of the brain include an interesting set of hand-motor control regions and is a source for rubrospinal as well as reticulospinal motor fibers (Alawieh et al., 2017; Gaubatz et al., 2020; Rüber et al., 2013, 2012). Thus, the possible activation of frontomesial motor regions with high tDCS dose in the bihemispheric montage might have important implications for studies examining the importance of frontomesial, crossed and uncrossed, motor systems in motor learning studies in healthy subjects as well as skill relearning studies in patients with a stroke.

### Relationship between normJ, rCBF and FSP

Although our experimental design and montage variation is relatively simple, results suggest that the effects of montage are difficult to predict and need to be examined both with biological data as well as modeling data and there could be a difference between the surrogate brain data and the simulated data. Electrical field models suggest that the unihemispheric montage primarily sends current from the targeted sensorimotor regions through contralateral premotor and ipsilateral prefrontal regions, whereas bihemispheric stimulation targets motor and premotor regions bilaterally. We had hypothesized that the ASL results would correlate with the simulated electrical current, however, there was no such correlation observed pointing out difficulty to predict dynamic biophysiological activity such as blood flow using finite element modeling. The correlation between finger sequence performance for each hand and simulated current density from the contralateral ROI showed strong correlation. Further, correlation between right hand finger sequence performance and simulated current density from the right hemispheric ROI was also significant. This correlation between finger sequence performance and simulated results help us estimate how well the model predicts behavioral results.

### Safety and Tolerability

We administered a total of 266 tDCS sessions in the laboratory and in the MR scanner; 94 of those were with 4mA stimulation intensity. We checked for the presence of adverse events including skin burns or other skin lesions as well as any abnormality in diffusion weighted imaging findings under the electrode during and immediately after tDCS session. No significant adverse effects occurred and the concurrent tDCS-MR imaging did not lead to any consistent image artifacts. Our results show that noninvasive brain stimulation with anodal tDCS up to 4mA is safe and tolerable, although tolerability scores showed a dose effect with higher doses leading to higher scores on the VAS. Our results align with the published literature suggesting that current intensity up to 4mA is safe and tolerable (Chhatbar et al., 2017; Dagan et al., 2018; Khadka et al., 2017; Trapp et al., 2019). Further, the current intensity of 4mA is smaller than the predicted lower threshold that might theoretically cause brain damage (Bikson et al., 2016).

With 94 tDCS sessions with 4mA applied current intensity, our results provide strong support for the safety and tolerability for high intensity tDCS. Analysis of the tolerability scores showed a significant dose effect. However, the correlation of the ‘tolerability score from behavioral experiment and left hand finger sequence performance’ and ‘tolerability score from tDCS-MR experiment and MR outcome measures’ were not significant. This suggests that the tDCS effects observed in our study cannot be explained by tolerability effects and are not explained by differential skin sensations experienced by our volunteers undergoing different tDCS-DOSE stimulation experiments.

### Limitations

One of the limitations of our study is that we did not record MEP data before and after tDCS sessions. MEPs have shown an effect of dose and stimulation duration. We opted to concentrate on a behavioral measure (e.g., FSP) and on a physiological measure (e.g., rCBF), which already required subjects to commit to 12 sessions. Another limitation of our study is incomplete data, since not every subject was able to finish 12 sessions: six different dose-montage combinations in a behavioral and six different dose-montage combinations in an imaging experiment. Using a linear mixed effects model allowed us to correct for missing data.

## Abbreviations

rCBF: regional cerebral blood flow
FSP: Finger Sequence Performance
tCD: Total Charge Density
VAS: Visual Analog Scale

## Acknowledgement

This research was supported by an NIH BrainInitiative grant (RO1MH111874). Dr. Schlaug also acknowledges support from U01NS102353. We are thankful for and very much appreciate the generous support that Klaus Schellhorn from neuroConn has provided to us with making a state-of-the-art multi-channel MR DC stimulator available to us and his helpful suggestions with setting up the concurrent tDCS-MR experiments and being available for any trouble shooting over the years. We would also like to thank Fotini Papadopoulou, our MR technologist, for her support with all of the complicated and lengthy concurrent tDCS-MR acquisitions.

## Notes

### Competing Interest Statement

The authors have declared no competing interest.

### Summary of Updates

1. Captions of Figure 1 and Figure 5 2. Discussion section

